# Electrostatics cause the molecular chaperone BiP to preferentially bind oligomerized states of a client protein

**DOI:** 10.1101/2021.10.11.463904

**Authors:** Judy L.M. Kotler, Wei-Shao Wei, Erin E. Deans, Timothy O. Street

## Abstract

Hsp70-family chaperones bind short monomeric peptides with a weak characteristic affinity in the low micromolar range, but can also bind some aggregates, fibrils, and amyloids, with low nanomolar affinity. While this differential affinity enables Hsp70 to preferentially target potentially toxic aggregates, it is unknown how Hsp70s differentiate between monomeric and oligomeric states of a target protein. Here we examine the interaction of BiP (the Hsp70 paralog in the endoplasmic reticulum) with proIGF2, the pro-protein form of IGF2 that includes a long and mostly disordered E-peptide region that promotes proIGF2 oligomerization. We discover that electrostatic attraction enables the negatively charged BiP to bind positively charged E-peptide oligomers with low nanomolar affinity. We identify the specific BiP binding sites on proIGF2, and although some are positively charged, as monomers they bind BiP with characteristically low affinity in the micromolar range. We conclude that electrostatics enable BiP to preferentially recognize oligomeric states of proIGF2. Electrostatic targeting of Hsp70 to aggregates may be broadly applicable, as all the currently-documented cases in which Hsp70 binds aggregates with high-affinity involve clients that are expected to be positively charged.

## Introduction

Hsp70-family chaperones are crucial molecular machines involved in folding nascent polypeptides, holding non-native state protein folding intermediates, and disaggregating misfolded proteins^1,2^. They bind exposed, extended, and hydrophobic segments of unfolded, misfolded, or partially-folded proteins^3,4^. Hsp70s are composed of two domains held together by an interdomain linker: a nucleotide-binding domain (NBD) and substrate-binding domain (SBD) (Figure 1A). The SBD contains a beta-sheet region (SBD_β_), including the hydrophobic substrate-binding cleft, and an alpha helical lid (SBD_α_). Hsp70s populate two major conformations that are dictated by the nucleotide bound in the NBD. In the ATP-bound conformation, the NBD and SBD_β_ are docked, the linker is bound to the NBD, and the SBD_α_ lid is open to expose the SBD_β_ substrate-binding cleft^5^. After ATP is hydrolyzed, the NBD and SBD_β_ undock, and the SBD_α_ lid closes onto the SBD_β_ substrate-binding cleft^6^. The ADP-bound conformation typically favors client binding, in which a client can be trapped between the SBD_β_ substrate-binding cleft and SBD_α_ lid^5^. BiP, like other Hsp70s, is negatively charged, much of which is contributed by the SBD (Figure 1A).

**Figure 1.**
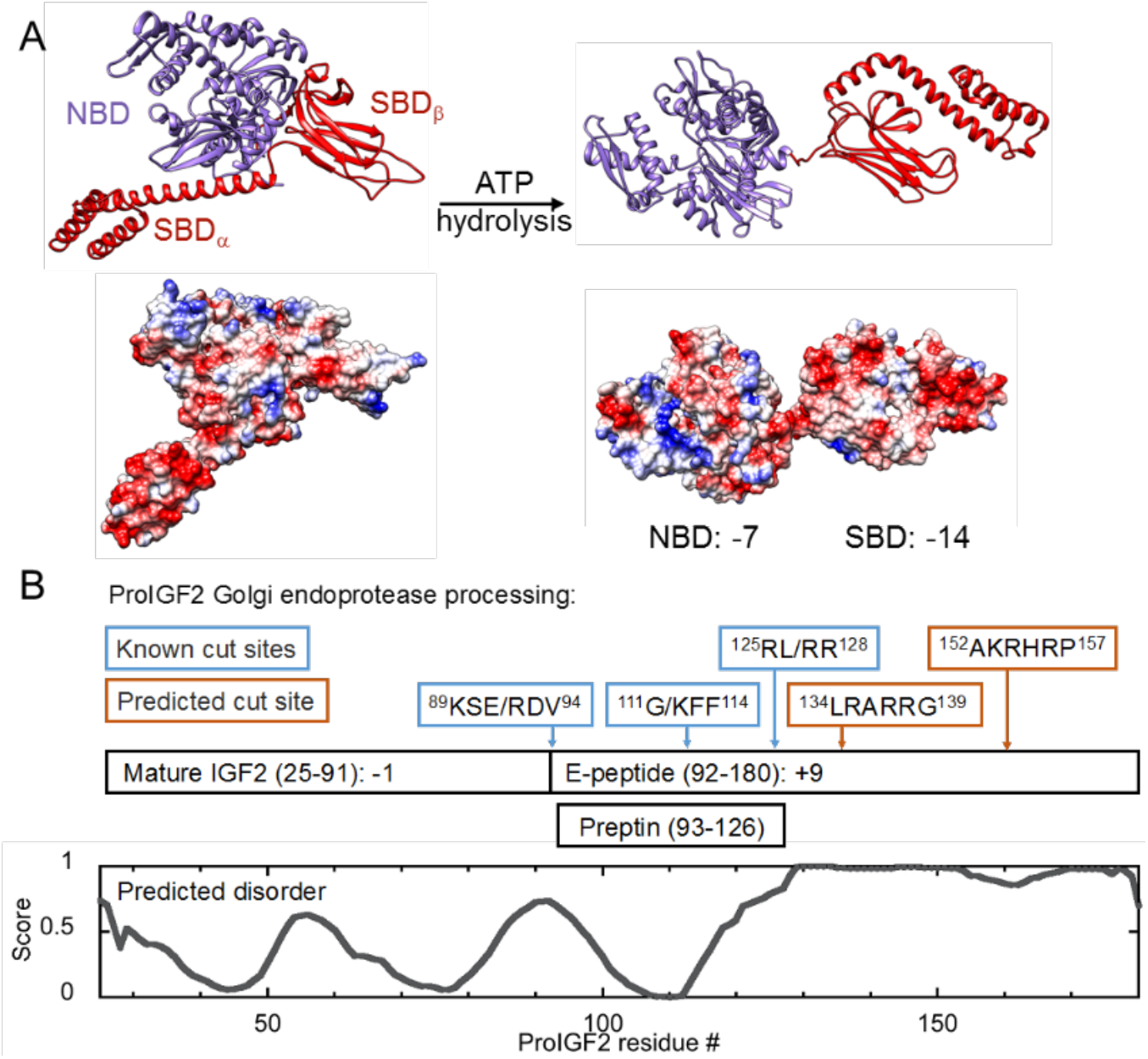
Overview of BiP and proIGF2. **(A)** BiP’s two major conformations are shown in ribbon and surface, with calculated electrostatic potential coloring (red for negative charge and blue for positive charge)^18^. BiP’s ATP-bound conformation has NBD and SBD docked (PDB:5E84), while BiP’s ADP-bound conformation has domains undocked (BiP homology model, PDB:2KHO)^5,6^. The net charges of the NBD and SBD are noted below the ADP conformation. **(B)** Known and predicted endoprotease processing sites on proIGF2. “/” indicates known cut site within sequence. Predicted sites are based on furin protease motif, “xBxBB/x”, where x is an uncharged and B is a basic (Arg or Lys) amino acid^19^. Amino acid sequences shown in boxes are from *Mus musculus* proIGF2. Disorder for proIGF2 amino acid sequence was predicted by PONDR^20^.

A range of neurodegenerative diseases are associated with the formation of protein aggregates and fibrils, and it is important to understand how Hsp70-type chaperones differentiate whether the “client” protein is in an oligomeric or monomeric state. In some cases Hsp70s bind clients with much higher affinity when the client is oligomeric. For example, human Hsp70 binds tau fibrils with low nanomolar affinity and tau monomers with micromolar affinity^7^. In contrast, human Hsp70 binds α-synuclein fibrils and monomers with comparable low micromolar affinities^8^. It is unknown how Hsp70 achieves high affinity for oligomeric client states, and why this affinity enhancement is observed for some but not all oligomeric clients.

ProIGF2 is the pro-protein of insulin-like growth factor (IGF) 2, which is a member of the insulin family of hormones, and is a mitogen for fetal and placental cell growth^9^. ProIGF2 is targeted to the ER via an N-terminal (24 residue) ER-signaling sequence that is cleaved upon entrance to the ER. Following the signal sequence is the 67 residue mature hormone region and 89 residue positively-charged E-peptide (Figure 1B). Folded, α-helical mature IGF2 contains three disulfide bonds, whereas the E-peptide is predicted to be mostly disordered and has minimal secondary structure^10^. Once folded, proIGF2 is translocated from the ER to the Golgi for further processing^11^. ProIGF2 is modified by N-acetylgalactosamine likely in the *cis*-Golgi, and sialic acid addition and oligosaccharide maturation in the *trans*-Golgi^11^. Modified proIGF2 is proteolytically cleaved twice by the proprotein convertase PC4: first to the intermediate form (residues 25-126) and then to mature IGF2 (25-91) (Figure 1B)^12,13^. The second cleavage liberates the hormone preptin (93-126)^14^. Preptin, which has minimal structure^15^, is cosecreted with insulin and amylin and increases glucose-mediated insulin secretion from pancreatic β-cells^16^. An intermediate cleavage product (25-111) has also been observed in bovine serum^17^. The positively-charged cleavage motifs confer a net charge of +9 to the E-peptide, while mature IGF2 has a net charge of −1.

Previous work demonstrated that proIGF2 forms dynamic oligomers, where the E-peptide region is necessary for oligomerization^10^. BiP and the ER Hsp90 paralog Grp94 regulate the assembly of these oligomers while exerting only a minimal influence on the folding of proIGF2^10^. It was left unknown where and how BiP and Grp94 interact with proIGF2. For example, whether BiP and Grp94 compete for binding sites on proIGF2 or whether they recognize different areas, and how tightly these chaperones interact with proIGF2 oligomers. Here, by dissecting the mechanism by which BiP recognizes proIGF2 oligomers, we discover that electrostatics play a defining role. Given the available data in the literature, electrostatics provide a plausible explanation of why Hsp70 chaperones preferentially bind some aggregated clients, such as tau, but not other clients such as α-synuclein.

## Results

### BiP binds E-peptide oligomers with high affinity

We first utilized dynamic light scattering (DLS) to quantify the size of proIGF2 and E-peptide oligomers and the range of conditions in which oligomers are formed. ProIGF2 oligomers are larger than E-peptide oligomers and in both cases their size increases with protein concentration (Supplemental Figure 1A). In these experiments proIGF2 was maintained in a reduced and non-native state by the reducing agent TCEP. We evaluated proIGF2 concentrations at 1μM and below, because at higher concentrations proIGF2 forms large particles that produce optical light scattering (Supplemental Figure 1B), which prevents accurate size determination by DLS. E-peptide oligomers and mature IGF2 do not scatter light at concentrations up to 5μM (Supplemental Figure 1B). For proIGF2, the build-up of light-scattering particles is slower at pH 6 versus at pH 7.5 (Supplemental Figures 1C,D), so the first experiments were performed at this lower pH condition.

Because the hydrodynamic radius (R_H_) of proIGF2 and E-peptide oligomers are in the range of hundreds of nanometers (Supplemental Figure 1A), much larger than BiP (R_H_ ∼ 3nm^10^), we reasoned that the binding of BiP to these oligomers could be measured by fluorescence depolarization (FP). Specifically, if BiP preferentially binds monomers or small oligomers then a negligible increase in polarization is expected, due to the small size of proIGF2 (17 kDa) relative to BiP (70 kDa), whereas if BiP preferentially binds large oligomers then a large change in polarization is anticipated (Figure 2A). Figure 2B shows that BiP binds both proIGF2 and E-peptide oligomers, whereas no FP change is observed for mature IGF2. BiP binding to proIGF2 and E-peptide oligomers is observed under both ATP (Figure 2B) and ADP (Supplemental Figure 2) conditions. The larger amplitude of FP change for proIGF2 versus E-peptide is consistent with the larger size of proIGF2 oligomers. In both cases the FP signal increases with protein concentration similar to the increasing size of E-peptide and proIGF2 oligomers as measured by DLS. BiP’s SBD is responsible for the high-affinity binding of proIGF2 oligomers because the isolated BiP NBD has only weak interactions with proIGF2 (Supplemental Figure 2).

**Figure 2.**
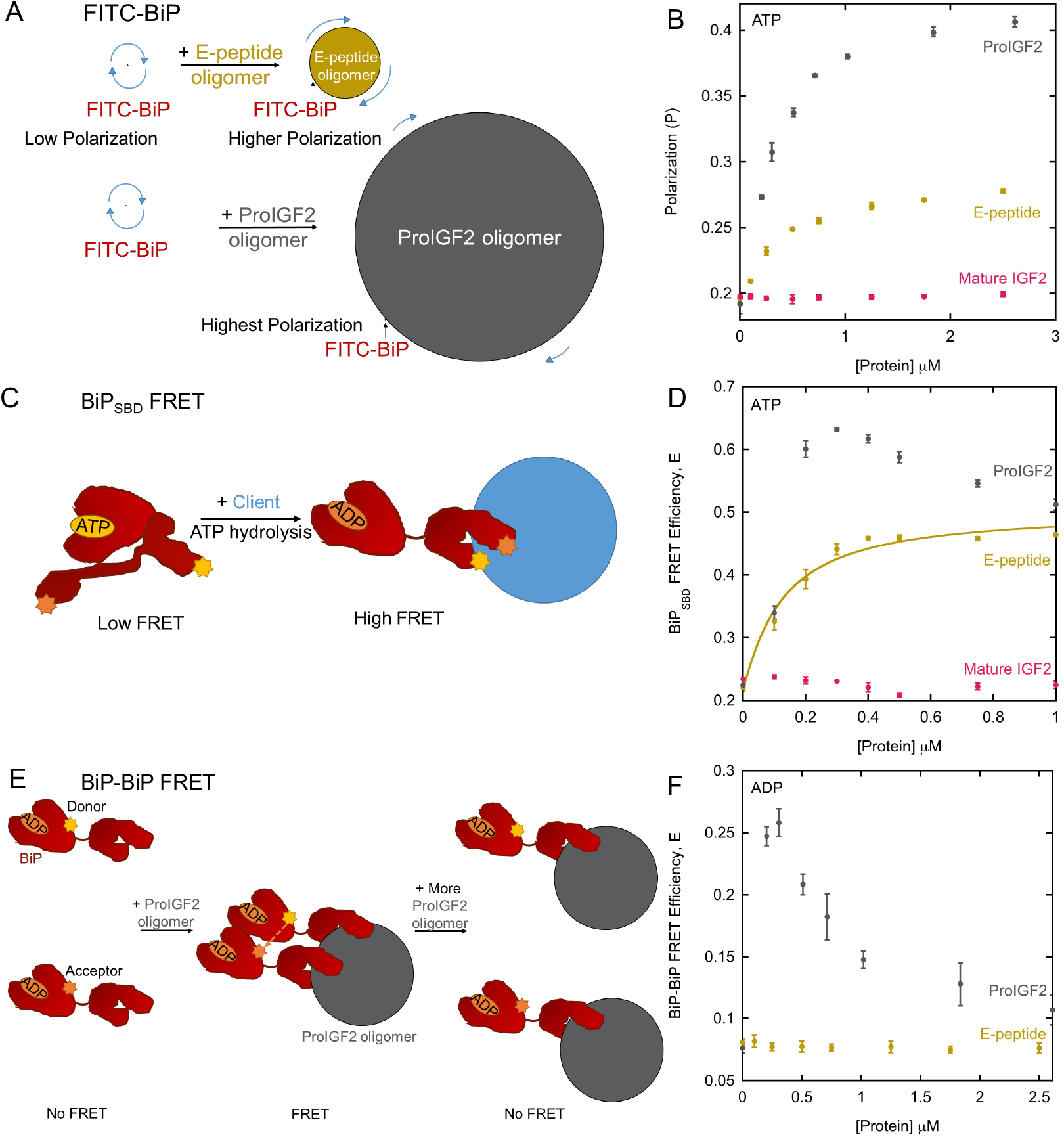
BiP binds proIGF2 and E-peptide oligomers. **(B)** Schematic of FITC-BiP fluorescence depolarization assay. BiP, E-peptide oligomer, and proIGF2 oligomer relative sizes are shown at 1 μM. Blue arrows indicate relative tumbling rates where longer arrows indicate faster tumbling and shorter arrows indicate slower tumbling. **(C)** FP assay with FITC-BiP and proIGF2, E-peptide, and mature IGF2 in the presence of ATP. **(D)** Schematic of BiP_SBD_ FRET where BiP’s lid-open, ATP conformation produces low FRET efficiency (E) and BiP’s lid-closed and client-bound, ADP-conformation produces high FRET efficiency. Donor and acceptor labels can be present in both locations on the SBD due to labeling protocol, but only one of each is shown for clarity. BiP and oligomer sizes are not to scale. **(E)** BiP_SBD_ FRET data for proIGF2, E-peptide and mature IGF2. Solid line is a fit to equation 4, with K_D_ = 0.098 ± 0.010 μM. **(F)** Schematic of BiP-BiP FRET experiment with BiP separately-labeled with either donor or acceptor fluorophore. BiP and oligomer sizes are not to scale. **(G)** BiP-BiP FRET data for proIGF2 and E-peptide. Error bars are the SEM for three replicates.

The above FP assay cannot yield a binding affinity because the FP signal is determined by the oligomer size, however, given that large FP changes are observed at sub-micromolar concentrations of proIGF2 and E-peptide, the FP data suggests sub-micromolar affinity. To determine BiP affinity for oligomers, we utilized a FRET assay measuring the conformation of BiP’s SBD^21,22^. This BiP_SBD_ FRET assay produces low FRET efficiency in the ATP-bound, lid-open state and high FRET efficiency in the ADP-bound and lid-closed state (Figure 2C). ProIGF2, E-peptide, and mature IGF2 were assayed with BiP_SBD_ FRET in the presence of ATP (Figure 2D). BiP_SBD_ FRET increases upon binding proIGF2 and the E-peptide oligomers, indicating lid closure, as is typically observed when an Hsp70 binds a peptide client^21,23^. No lid-closure is observed in the presence of mature IGF2. The FRET change for E-peptide can be fit with a binding curve (Figure 2D, solid line) yielding a binding affinity of approximately 100 nM, approximately 100-fold higher affinity than is typical for BiP binding a monomeric peptide under ATP conditions. While measuring BiP’s conformation is an indirect determination of binding affinity, later we demonstrate that this indirect method agrees with BiP affinities measured directly for peptides using an FP assay.

The lid closure of BiP in the presence of E-peptide oligomers could result from stable binding while the BiP ATPase cycle is stalled in the ADP state, or from BiP cycling through rounds of ATP binding and hydrolysis with accelerated ATPase kinetics that shift the conformational equilibrium towards the lid closed state. ATPase measurements support the latter case, as the BiP ATPase rate increases from 0.23 ± 0.01 min^-1^ to 1.53 ± 0.02 min^-1^ in the presence of 2.5 μM E-peptide. Due to the enhanced hydrolysis of ATP by BiP, the measured affinity under ATP conditions will have a contribution from the ADP-bound state, and we sought to measure this contribution. However, measuring BiP affinity under ADP conditions is challenging because BiP is maintained uniformly in the high-FRET lid-closed state, so no change in FRET efficiency is observed (Supplemental Figure 3). Therefore, we utilize ADP conditions with trace quantities of ATP, to enable a change of FRET to be measured. For example, commercial stocks of ADP contain ∼2% ATP (see Figure 1 in Liu et al.^24^), which we remove by a pretreatment with hexokinase (HK, see Methods). In experiments with this residual ATP present (termed “ADP, no HK”) or with an additional 5% added ATP, we can measure BiP affinity to E-peptide oligomers under predominantly ADP conditions. In both cases, the measured BiP affinity to E-peptide oligomers is in the range of 10-20 nM (Supplemental Figure 3). The roughly ten-fold higher affinity of BiP for E-peptide oligomers under ADP versus ATP conditions is similar to the nucleotide dependence observed for other Hsp70s binding peptides^25,26^.

**Figure 3.**
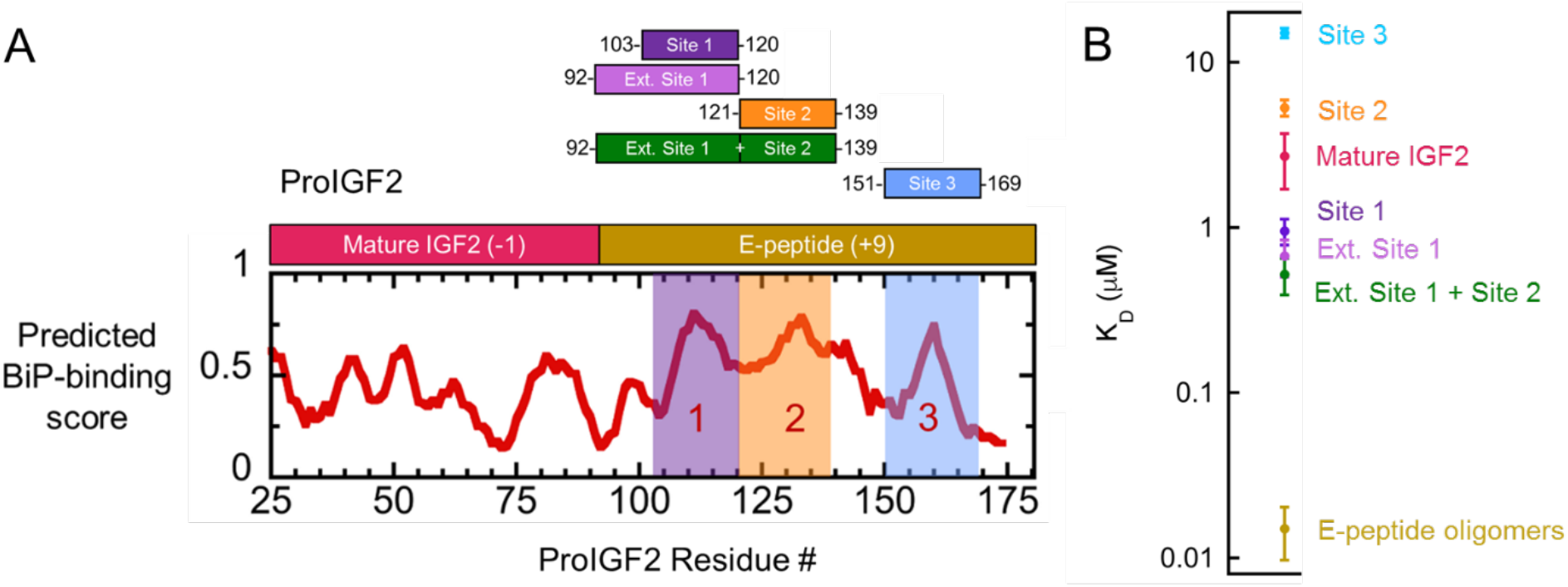
**(A)** BiP-binding sites (labeled 1, 2, and 3) as predicted by BiPPred^27^. Color shading indicates FITC-labeled peptide binding sites 1 (purple), 2 (orange), and 3 (blue). **(B)** Affinities of monomeric E-peptide fragments (from Supplemental Figure 4) and the oligomeric E-peptide (from Supplemental Figure 3). Measurements performed under ADP conditions. Error bars are the SEM for at least three binding curve replicates.

Unlike the BiP_SBD_ FRET data with E-peptide oligomers in Figure 2D, which can be fit to a binding curve, for proIGF2 oligomers the FRET efficiency first rises above 0.5 and then falls back to a saturating value close to that observed for the E-peptide. Due to the fluorophore labeling scheme (Methods) the maximum FRET efficiency is 0.5 for a BiP monomer. However, if BiP monomers are positioned closely on an oligomer then FRET efficiencies above 0.5 could arise from an additional contribution from FRET between BiPs. We developed a FRET assay to detect when BiPs are in close proximity (“BiP-BiP FRET”, Figure 2E). Upon adding proIGF2, BiP-BiP FRET reaches a maximum at 0.3 μM proIGF2, indicating multiple BiP’s are occupying a single proIGF2 oligomer. Higher concentrations of proIGF2 decrease FRET (Figure 2F). At these higher concentrations of proIGF2 oligomers, with the same concentration of BiP, single BiPs will occupy different proIGF2 oligomers and FRET efficiency will decrease. Multiple BiPs binding per oligomer is not necessary for high affinity, however, because BiP-BiP FRET is not observed in experiments with E-peptide (Figure 2F).

Overall, we conclude that E-peptide oligomers are well-suited to uncover the origin of BiP’s high affinity for oligomers. Unlike proIGF2, E-peptide oligomers are not confounded by BiP-BiP FRET, making the BiP_SBD_ FRET assay a powerful tool for measuring BiP affinity to oligomers. Furthermore, whereas proIGF2 experiments are performed at low pH to limit the formation of very large oligomers that scatter light, E-peptide oligomers are well behaved at both low and high pH values (Supplemental Figure 1B) and BiP binds with comparable affinity at both pH 6.0 (K_D_ of 98 ± 10 nM) and pH 7.5 (K_D_ of 130 ± 10 nM) under ATP conditions.

### Identification of BiP binding sites on proIGF2

BiP binding sites can be predicted from primary sequence^27^ and three potential sites are on the E-peptide (Figure 3A, labeled 1, 2, and 3). Binding site 1 resides within the preptin region. We also evaluated BiP’s interaction with the mature region of proIGF2. All E-peptide peptide constructs were labeled with FITC for FP measurements (Methods) and maintained at a low concentration (50 nM) in BiP binding experiments to suppress oligomerization. Indeed, in the absence of BiP sites 1-3 all have similar low polarization values of ∼0.07 (Supplemental Figures 4A-C) that are characteristic of monomeric peptides.

**Figure 4.**
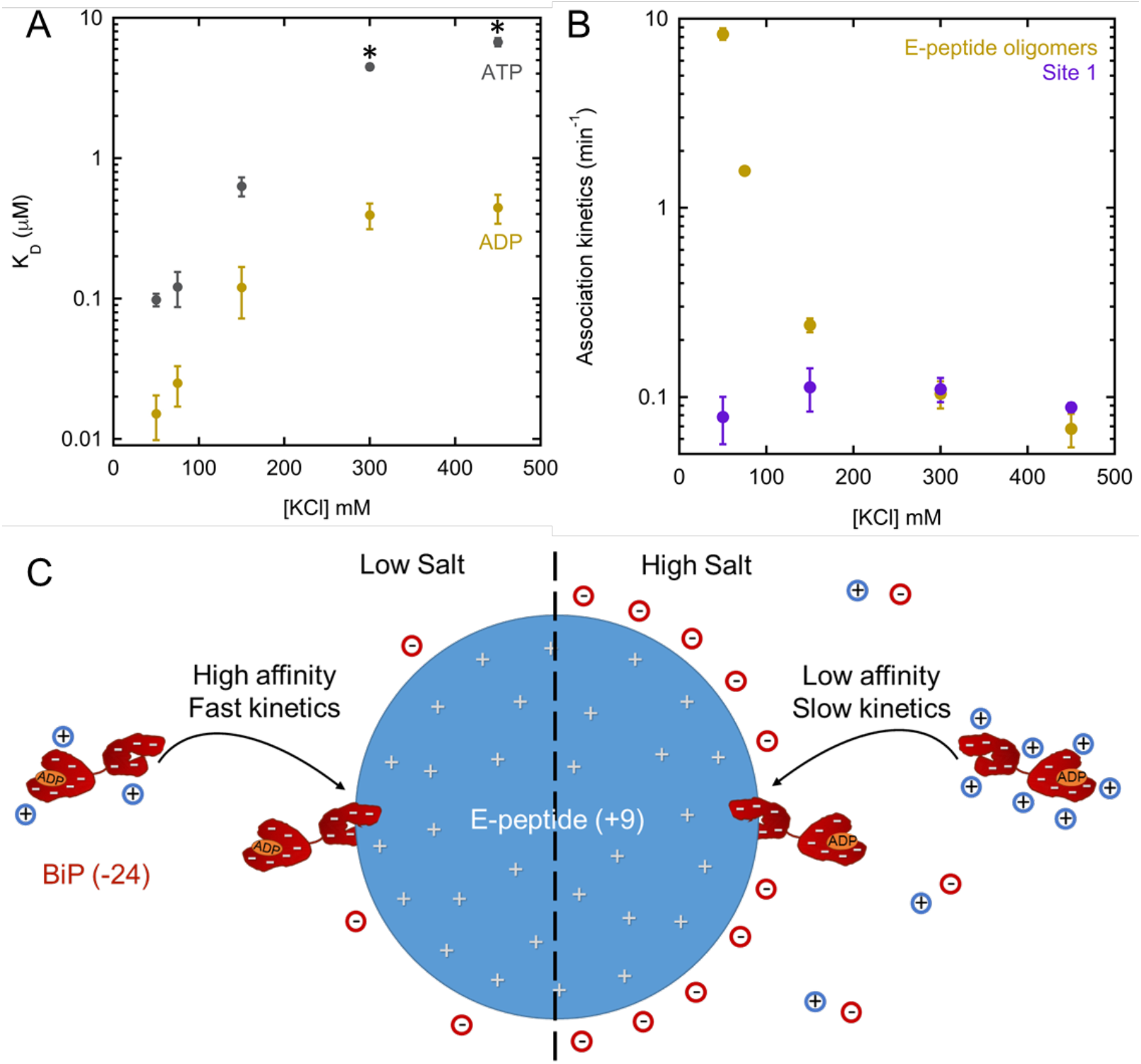
Influence of salt on BiP binding E-peptide oligomers. **(A)** Salt dependence of K_D_ for BiP binding E-peptide as measured by BiP_SBD_ FRET assay in the presence of ADP or ATP. Individual binding curves shown in Supplemental Figure 8A&B. Asterisks indicate lower confidence of fitting, as described in Supplemental Figure 8B. **(B)** Salt dependence of association kinetics between BiP and 0.1 μM E-peptide as measured by BiP_SBD_ FRET assay under ADP conditions. Association kinetics between site 1 and 0.1 μM BiP was determined by FP (Methods). **(C)** Model for BiP’s salt-dependent binding of E-peptide oligomers. Negative charge for BiP and positive charge for E-peptide is shown as grey dashes and grey plus-signs, respectively. Counterions to E-peptide and BiP are shown as negatively-charged red circles and positively-charged blue circles, respectively.

BiP binds all three binding-site peptides and mature IGF2 with low micromolar affinity, in the presence of ADP, and approximately 10-fold weaker affinity in the presence of ATP (Supplemental Figures 4A-D). Site 1 has the highest affinity for BiP (K_D_ ∼ 1μM under ADP conditions), a much weaker binding than is observed for E-peptide oligomers (K_D_ ∼ 10-20 nM). While this difference in affinity could plausibly be explained if site 1 is not a complete BiP binding site, we confirmed that site 1 is complete by constructing E-peptide fragments centered at site 1, that are extended in the N-terminal direction (residues 92-120) and in the C-terminal direction (residues 92-139). Both “extended fragments” and site 1 bind BiP with similar affinity under ADP conditions (compare Supplemental Figures 4A, E, and F). Because the 92-139 fragment contains both site 1 and site 2, we can exclude the possibility that the high affinity of BiP to E-peptide oligomers is due to an avidity effect from these two closely spaced BiP binding sites. This is consistent with the absence of BiP-BiP FRET on the E-peptide (Figure 2F). Site 1 can outcompete site 2 in binding to BiP under ADP conditions, with characteristically slow displacement kinetics (Supplemental Figure 5A). We conclude that site 1 is the dominant BiP binding site on proIGF2.

Grp94 has minimal binding for sites 1-3 and mature IGF2 under both ATP and ADP conditions (Supplemental Figure 6), demonstrating that these sites are specific to BiP. Interestingly, site 1 binds non-specifically to BSA whereas sites 2&3 do not bind BSA (Supplemental Figure 7), suggesting that BiP’s preferential binding to site 1 may serve a biological role in preventing non-specific interactions with this region of the E-peptide. We utilized the slow displacement kinetics of site 2 to test whether BiP binds E-peptide oligomers specifically. If BiP binds E-peptide oligomers specifically, a BiP:site 2 complex must first release site 2 before binding the E-peptide oligomer, and the displacement kinetics should be slow. If BiP binds E-peptide oligomers non-specifically no such displacement will be observed. E-peptide oligomers show slow displacement kinetics that are similar to that of site1 and site 2 (Supplemental Figure 5B) indicating that BiP binds E-peptide oligomers specifically and in a manner similar to a typical peptide client, albeit with much higher affinity.

### Electrostatic steering enhances BiP affinity for E-peptide oligomers

The charge difference between BiP and both proIGF2 and the E-peptide (Figure 1) suggests that high affinity binding might have an electrostatic contribution. If true, BiP affinity for proIGF2 and E-peptide oligomers should be salt dependent due to charge screening. The affinity of BiP for E-peptide oligomers as measured by BiP_SBD_ FRET is indeed highly salt dependent, where increasing the salt concentration weakens BiP’s affinity for E-peptide oligomers under both ADP and ATP conditions (Figure 4A, Supplemental Figures 8A-B). The highest salt concentration data in the presence of ATP requires fixing the saturating FRET efficiency value, and therefore these K_D_ values are not as well defined and should be interpreted cautiously (these data are marked with an asterisk in Figure 4A). While salt-dependent affinities cannot be measured for BiP and proIGF2 because of BiP-BiP FRET (Figures 2D,F), the FP assay described in Figure 2A shows a loss of binding between BiP and proIGF2 oligomers with increasing salt (Supplemental Figure 8C).

The strong salt dependence of BiP binding E-peptide oligomers is observed with different salts (KCl, NaCl, and KI, see Table 1), as expected for electrostatic screening rather than a specific ionic interaction. ProIGF2 light scattering is minimally salt-dependent (Supplemental Figures 1C,D), and E-peptide does not scatter light at any salt condition tested (Supplemental Figure 1C), suggesting that oligomer size changes cannot explain the strong salt-dependent biding of BiP to both proIGF2 and E-peptide.

**Table 1.** BiP dissociation constants for E-peptide using BiP_SBD_ FRET assay with different salts, NaCl, KI, and KCl, in the presence of ADP.

| [Salt] mM | NaCl, $K_D$ (μM) | KI, $K_D$ (μM) | KCl, $K_D$ (μM) |
| --- | --- | --- | --- |
| 50 | $0.0047 \pm 0.0022$ | $0.021 \pm 0.003$ | $0.015 \pm 0.005$ |
| 300 | $0.29 \pm 0.03$ | $0.51 \pm 0.10$ | $0.39 \pm 0.08$ |

Given the dramatic influence of salt on the affinity of BiP to E-peptide oligomers, we questioned whether electrostatic steering, which is characterized by salt-dependent association rates leading to salt-dependent binding affinity, is the underlying cause. At low salt BiP binding is indeed accelerated and we therefore performed the measurements at 10°C to slow the kinetics such that they can be quantified over a large range of rates. Figure 4B shows salt-dependent association rates between BiP and E-peptide in which association kinetics decrease by ∼100-fold between 50 to 450 mM KCl. In comparison, Site 1 has no salt-dependent changes in association rates and converges with E-peptide association rates at high salt. Collectively, the above results indicate that electrostatic steering enhances BiP affinity for proIGF2 and E-peptide oligomers (Figure 4C).

### Two energetic contributions to BiP binding to E-peptide oligomers

Comparing the salt-dependent affinity of BiP to E-peptide oligomers versus its affinity for monomeric peptides (Figure 5A) shows that two distinct energetic contributions underpin BiP’s high affinity for E-peptide oligomers. BiP binds sites 1-3 with minimal salt-dependence, even though binding sites 2 and 3 are positively charged. This lack of salt dependence is also observed for the extended fragments centered at site 1 (residues 92-120 and 92-139). Importantly, BiP’s affinity for E-peptide oligomers at high salt matches that of all the monomeric fragments that include site 1. This suggest that two energetic contributions enable BiP to bind the E-peptide oligomers with high affinity. The first contribution is from the typical binding affinity between BiP and site 1, and the second contribution is from electrostatic attraction between BiP and the E-peptide oligomer. As the electrostatic contribution is screened by salt BiP’s affinity for E-peptide oligomers converges to the measured affinities of all constructs that contain site 1.

**Figure 5.**
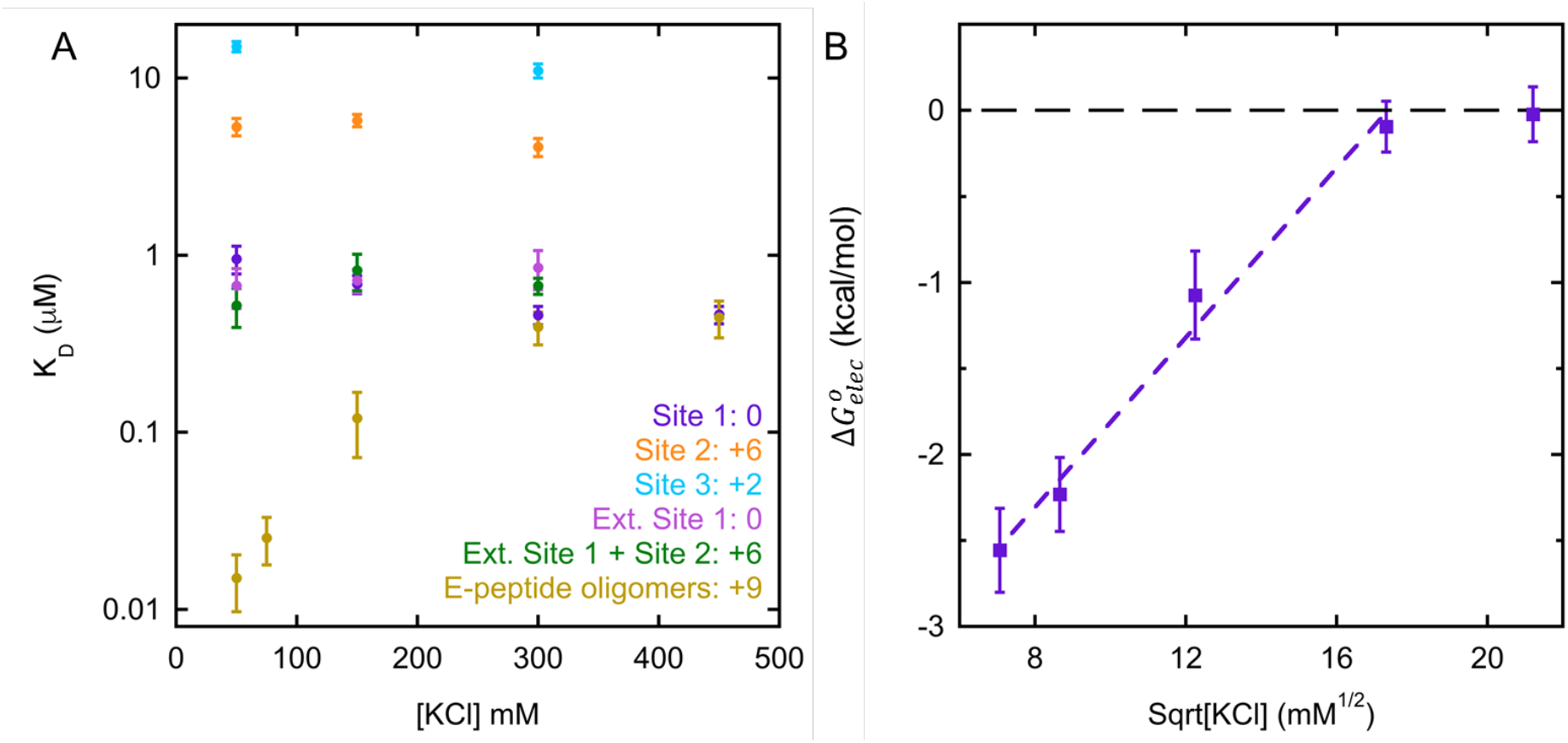
Compilation of BiP’s affinity for E-peptide and binding sites in presence of increasing salt concentrations. **(A)** Affinities of monomeric E-peptide fragments and the oligomeric E-peptide. Net charge of each fragment is indicated next to name. Error bars are the SEM for at least three binding curve replicates. **(B)** Electrostatic contribution as calculated by 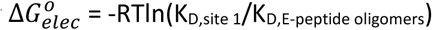, which is screened out by increasing salt^28^. Error bars are the propagated uncertainty from panel A. The K_D,site 1_ value at 50mM KCl is used to calculate 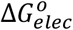 at both 50 and 75 mM KCl. Purple dashed line indicates linear fit (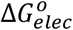 intercept: −4.3 ± 0.2 kcal/mol and slope: 0.25 ± 0.02). Black dashed line indicates 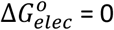.

The data in Figure 5A enables the electrostatic contribution to binding to be determined at any salt concentration. For example, under ADP conditions and at 50 mM KCl BiP binds E-peptide oligomers with ∼60-fold higher affinity than to site 1 (K_D_ = 0.015 ± 0.005 μM compared to K_D_ = 0.95 ± 0.17 μM). A similar analysis under ATP conditions shows that electrostatic attraction provides a ∼130-fold enhancement of affinity of BiP to E-peptide oligomers at 50 mM KCl (Supplemental Figure 9). While the influence of salt has a dramatic influence on BiP binding oligomers, only minor effects are observed for BiP binding the monomeric site 1 peptide. For example, increasing salt provides a slight enhancement of site 1 binding under ADP conditions (K_D_ of 0.95 ± 0.17 μM at 50 mM KCl versus 0.46 ± 0.05 μM at 450 mM KCl), and no salt-dependent affinity changes are observed under ATP conditions (Supplemental Figure 9). The agreement between the affinities measured indirectly using BiP_SBD_ FRET assay and directly using FP at high salt confirms that the BiP_SBD_ FRET measurements provide a reliable measurement of client affinity.

Theoretical predictions (see Chapter 9 in Physical Biology Of The Cell^28^) and experiments on viral capsids^29^ provide a quantitative framework for understanding the influence of salt on electrostatic screening around large macromolecular assemblies. In particular, the free energy contribution of electrostatics from charged spherical assemblies should vary linearly with the square root of the salt concentration. Figure 5B shows the electrostatic contribution to BiP binding free energy of BiP versus the square root of the salt concentration. We indeed observe this expected linear relationship up until the point at which the electrostatics no longer contribute to binding (when 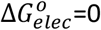). While other factors may also contribute to the binding of BiP to E-peptide oligomers, our data is consistent with a model in which two energetic factors predominate. The first is the foundational hydrophobic interactions between BiP and site 1, which has the characteristically weak affinity of ∼1μM and a minimal salt dependence. The second is the electrostatic contribution that is unique to the oligomerized the E-peptide and obeys the strong salt dependence predicted by electrostatic screening.

### The electrostatic driving force that favors BiP binding E-peptide oligomers is widely dispersed across the E-peptide sequence

E-peptide oligomers provide a favorable electrostatic contribution for BiP biding despite the fact that primary binding site on the E-peptide (site 1) has no net-charge. Thus, we expect that E-peptide variants containing site 1, but with lower net charge will have lower affinity for BiP. Recall that the E-peptide has clusters of positively charged residues at conserved endoprotease cut sites (Figure 1B). Therefore, we designed a series of truncations to successively remove each of these +3 charge clusters to determine whether a single charge cluster dominates or whether each cluster contributes equally (Figure 6A). In these experiments BiP is maintained at a low concentration, and the E-peptide constructs are titrated to enable oligomerization. Figure 6B shows a progressive enhancement of BiP affinity for E-peptide oligomers that are assembled from E-peptide constructs with progressively positive net charge. This trend (dashed lines, Figure 6B) shows a convergence of the high affinity binding of BiP to E-peptide oligomers to low affinity site 1 binding as the net charge on E-peptide constructs is reduced. This convergence is conceptually similar to the convergence in oligomer and site 1 affinities with increasing salt (Figure 6B). In both cases, negatively-charged BiP’s electrostatic targeting towards positively-charged E-peptide oligomers is screened physically by removing net charge from the E-peptide by way of truncations or by increasing the salt concentration.

**Figure 6.**
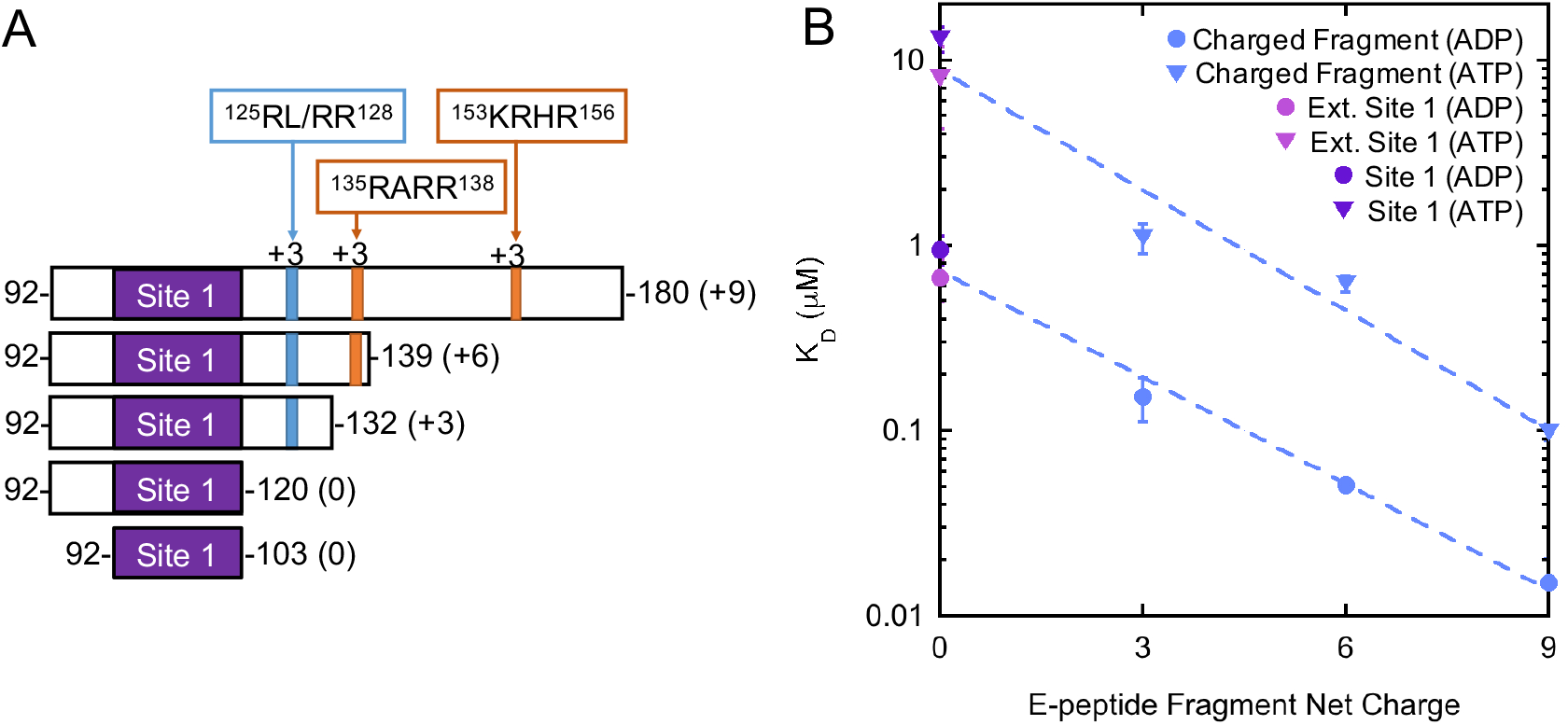
BiP’s interactions with E-peptide and E-peptide truncations of decreasing net charge. **(A)** E-peptide and E-peptide truncations with net charge and endoprotease cut sites are indicated for each construct. **(B)** BiP’s K_D_ for each E-peptide fragment listed in A, as a function of fragment’s net charge. K_D_ data is from BiP_SBD_ FRET, except for Ext. Site 1 and Site 1 which are from FP data. Error bars indicate SEM for 3 replicates, and error bars may be smaller than data point. Dashed lines indicate a logarithmic fit (K_D_ intercepts: 0.73 ± 0.10 μM (ADP); 8.9 ± 2.6 μM (ATP)). Slopes are: 10^−(0.19 ± 0.01)*x*^ (ADP); 10^−(0.22 ± 0.03)*x*^ (ATP), where *x* is the net charge of E-peptide fragment and the uncertainty is from the fitting error.

## Discussion

The action of Hsp70-type chaperones on aggregates, oligomers, and fibrils is a crucial aspect of cellular homeostasis, the adaptive response to environmental stress, and the progression of age-related diseases^30^. However, heterogeneity of aggregates, oligomers, and fibrils imposes technical challenges in determining how Hsp70s recognize oligomeric client states versus the monomeric peptide fragments that are often used as model systems to study Hsp70:client binding. Here, by dissecting the mechanism by which BiP recognizes proIGF2 and E-peptide oligomers, we discover that electrostatic attraction is a powerful driving force that enables BiP to preferentially bind oligomeric client states. BiP binds E-peptide oligomers with nanomolar affinity, but binds the monomeric constituent peptides with micromolar affinity. In this regard BiP interacts very differently with proIGF2 oligomers compared to the well-studied C_H_1 domain in which BiP binds full-length C_H_1 (K_D_ of 4.2 μM) with similar affinity to its constituent peptide motif (K_D_ of 12 μM)^21,31^. The predominant BiP binding site on proIGF2 is located at the preptin hormone region of the E-peptide (site 1, Figure 3A), a region that does not fold^32^. This again contrasts with the BiP recognition of C_H_1, in which the BiP binding site is buried after C_H_1 folds and forms a disulfide-linked complex with C_L_^31^. The preptin region binds non-specifically to BSA (Supplemental Figure 7), suggesting a functional role for BiP in protecting this region from non-productive interactions in the ER.

A strong electrostatic driving force causes BiP to preferentially bind oligomers. This is evident in the salt-dependent affinity of BiP to E-peptide oligomers but not to the constituent monomeric peptides (Figure 5A). The energetic contribution from electrostatic screening varies with the square root of the salt concentration (Figure 5B), as predicted theoretically^28^, and measured experimentally in the assembly of viral capsids^29^. The electrostatic affinity enhancement, spanning approximately two orders of magnitude, is reflected in the weakening influence of successive truncations that remove charge from the E-peptide (Figure 6B), and the salt-dependent association rate of BiP binding E-peptide oligomers (Figure 4B). Such salt-dependent association kinetics are characteristic of electrostatic steering between large highly charged complexes as with the positively charged multimeric Von Willebrand factor binding its negatively charged receptor glycoprotein Ibα^33^. In this case, the association kinetics span approximately two orders of magnitude between 80 to 500 mM salt^33^, a similar magnitude as what we observe for BiP binding E-peptide oligomers (Figure 4B).

The electrostatic explanation underlying BiP’s high affinity for oligomers provides a plausible explanation for why Hsp70 chaperones interact with certain aggregated client proteins with high affinity. Figure 7 is a compilation of previously measured Hsp70 affinities for monomeric and oligomeric clients, evaluated by the predicted net charge of the client (data and references are in Supplemental Table 1). Hsp70s bind peptides with a maximal affinity of ∼1 μM. Even engineered peptides that are designed to have high affinity for Hsp70, such as the Javelin sequence, only reach ∼1 μM^34^. A similar upper limit appears to apply to Hsp70 binding negatively charged clients such as α-synuclein (net charge of −9) irrespective of whether it is monomeric or oligomeric (K_D_ ∼10 μM)^8^. Clathrin is a second example of a negatively charged (net charge −64) oligomer, which Hsp70 also binds with low affinity (K_D_ ∼3 μM)^26^.

**Figure 7.**
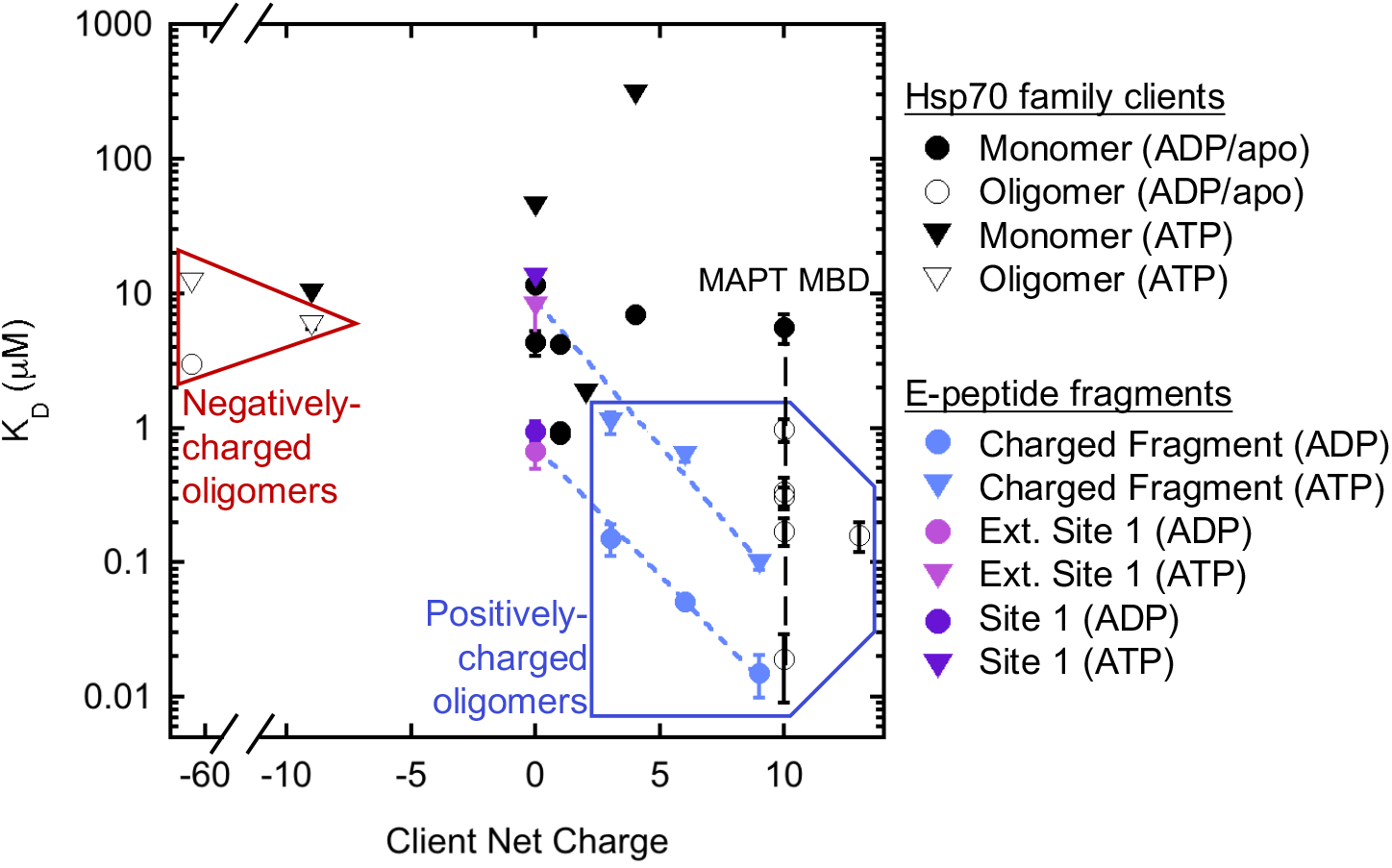
Compilation of Hsp70 and BiP dissociation constants to monomeric and oligomeric clients, sorted by net charge. See Supplemental Table 1 for data and references. Black dashed line indicates MAPT MBD K_D_ values with increasing size (1-2mer to fibril^7^). K_D_ data for E-peptide, E-peptide truncations, and associated blue dashed lines are from Figure 6.

To our knowledge the only reported instances of Hsp70 binding with much higher affinity than ∼1 μM is in the case of aggregates of positively charged clients. For example, the binding of cytosolic Hsp70 (net charge −11) to MAPT MBD (net charge +10) is directly proportional to the size of the oligomer size, with larger oligomers yielding higher affinities (Figure 7, black dashed line). Hsp70 binds MAPT MBD fibrils with ∼10 nM affinity^7^, a value comparable to BiP affinity to E-peptide oligomers. A fast association rate is observed between Hsp70 and MAPT MBD^7^ raising the possibility that electrostatic steering is at work, similar to our findings with BiP and E-peptide oligomers (Figure 4B). The affinity of BiP to E-peptide oligomers, as well as E-peptide truncations with different net charges (blue diagonal dashed lines, Figure 7) all fall within the range of values measured for other positively charged clients. The data in Figure 7 is restricted to metazoan Hsp70s, however experiments with the bacterial Hsp70 homolog DnaK (net charge −30) suggest that this mechanism also applies. Specifically, DnaK has been reported to bind positively charged IAPP oligomers in the pM-range^35^. However, the affinity of DnaK for oligomers may not be directly comparable to metazoan Hsp70 data in Figure 7, because DnaK can bind monomeric peptides with ten-fold higher affinity than is observed for metazoan Hsp70s (Supplemental Table 2).

The idea that electrostatics target Hsp70s to positively charged oligomeric clients, provides predictions for future experiments. For example, the MAPT 3R and 4R isoforms, which also have a positive net charge, exhibit high affinity binding to Hsp70 (Supplemental Table 1) but their oligomerization state has not been determined^36^. The electrostatic explanation predicts that MAPT 3R and 4R isoforms should be oligomeric. In the case of IGF proteins, electrostatics should favor the binding of BiP to oligomeric states of proIGF1 (net charge +18), but not proinsulin (net charge −3). One additional area for future investigation is to better understand the role of oligomer size heterogeneity on BiP affinity. While Hsp70 affinity for the MAPT MBD is proportional to the oligomer size (Figure 7, black dashed line) more detailed measurements are needed to determine if a similar relationship holds for BiP binding to E-peptide and proIGF2 oligomers.

Our findings have implications for how the ER responds to stress. A current model for the unfolded protein response (UPR) activation involves BiP binding to the luminal portion of key transmembrane proteins (IRE1 and PERK) that are held inactive when BiP is bound^37^. In this model, when the concentration of unfolded protein gets sufficiently high BiP will favor binding the client proteins rather than the UPR transmembrane proteins. Our results suggests that oligomerized clients within the ER could displace BiP from the UPR receptors due to the relatively high affinity of BiP towards oligomers. Thus, our results suggest that the UPR may be initiated by protein oligomerization/aggregation that could be independent of a rise in the concentration of unfolded proteins.

## Methods

### Bioinformatics

Predicted BiP-binding motifs on proIGF2 were calculated with BiPPred^27^. BiPPred calculates a predicted BiP-binding score for a 7-residue motif, and an average BiPPred score for each residue is calculated and plotted in Figure 3A. Net charge is calculated from sum of –(Asp + Glu) + (Lys + Arg) residues.

### Protein purification

6-his tagged BiP was purified via Ni-NTA affinity chromatography, and 6-his tag was cleaved with TEV. Subsequent Ni-NTA affinity chromatography removed 6-his tag and TEV, anion-exchange chromatography removed nucleotide bound to BiP, and BiP was buffer exchanged with size-exclusion chromatography. BiP was stored in 25 mM Tris pH 7.5, 50 mM KCl, 1 mM DTT, and 2% glycerol.

ProIGF2, E-peptide, E-peptide 92-139, E-peptide 92-120, and mature IGF2 were purified from inclusion bodies. E-peptide, E-peptide 92-139, and E-peptide 92-120 contained an N-terminal 6-histidine tag and cysteine mutation at Ser95 for FITC labeling. Briefly, inclusion bodies were washed and insoluble protein was denatured in an 8 M urea, 25 mM Tris buffer containing reducing agent TCEP. Protein was purified by ion-exchange chromatography and/or Ni-NTA affinity chromatography in denaturing conditions. Proteins used in FP assays were labeled with FITC-maleimide. Proteins were stored denatured in buffer containing 8 M urea.

BiP-binding sites 1 and 3 were synthesized by Alan Scientific (Gaithersburg, MD) and site 2 was synthesized by Genscript (Piscataway, NJ). All peptides are N-terminally labeled with FITC via an amino hexanoic acid linker. For FITC-Mature-1cys, mature IGF2 was mutated to remove all cysteines except C70, which was labeled with FITC.

### Fluorescence depolarization

50 nM FITC-labeled BiP D27C or BiP NBD D27C was incubated with buffer containing 25 mM MES pH 6.0, 50/150/300 mM KCl, 1 mM MgCl_2_, 1 mM nucleotide (ADP or ATP), 0.5 mg/mL BSA, and 1 mM DTT until polarization values reached equilibrium, for about 30 minutes, at 37°C. Experiments were also conducted in the absence of BSA, when noted. Clients were added directly from purified 8 M urea stock, except proIGF2 (diluted out of denaturant 1:10 in 50 mM MES pH 6.0, 2 mM TCEP and incubated 20-30 minutes). Fluorometer setup had an excitation wavelength at 492 nm and emission wavelength at 520 nm with 6 nm slit widths, and 1 second integration time.

FP experiments containing FITC-labeled BiP-binding site peptides used 50 nM of labeled peptides, except for FITC-Mature-1cys (57.5 nM). For ATP experiments, FITC-labeled peptide was added to BiP pre-incubated for 20 minutes in a buffer containing 25 mM buffer (MES or Tris), 50/150/300 mM KCl, 1 mM MgCl_2_, 1 mM ATP, 0.5 mg/mL BSA, and 1 mM DTT at 37°C. For ADP experiments, contaminating amounts of ATP were removed from 1 mM ADP with 0.005 units hexokinase, 1 mM glucose, and 5 mM MgCl_2_ and incubated 1 hour at 37°C. Polarization measurements for FITC-E-peptide 121-139 were taken with excitation at 493 nm, emission at 522 nm, 5 nm slit widths, and a 1 second integration time. Polarization measurements with FITC-E-peptide 103-120 and FITC-E-peptide 151-169 used 493 nm excitation wavelength, 518 nm emission wavelength, and 6 nm slit widths. FP experiments with FITC-E-peptide 92-120 or FITC-E-peptide 92-139 had an excitation wavelength of 492 nm and emission wavelength of 522 nm.

*K*_*D*_ values were calculated using the single-site binding equation,

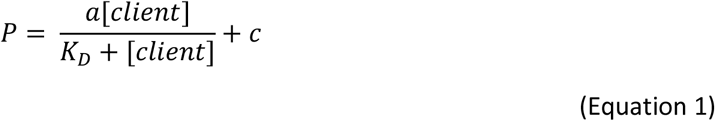

where *P* is polarization, *a* is the polarization amplitude and *c* is the polarization value in the absence of client. Association kinetics for 0.1μM BiP to site 1 (Figure 4B) was determined by a linear extrapolation of association kinetics measured over the complete range of BiP concentrations from Supplemental Figure 4A.

### FRET

For the BiP-BiP FRET assay, in separate reactions, BiP was labeled with donor (AlexaFluor 555 C_2_ maleimide) or acceptor (AlexaFluor 647 C_2_ maleimide) fluorophores. 25 nM donor-labeled BiP and 25 nM acceptor-labeled BiP were incubated until FRET efficiency reached equilibrium, about 20-30 minutes, in buffer containing 25 mM MES pH 6.0, 50 mM KCl, 1 mM ADP, 0.5 mg/mL BSA, and 1 mM DTT at 37°C. ADP was pretreated with 0.005 units hexokinase, 1 mM glucose, and 5 mM MgCl_2_ for 1 hour at 37°C. Clients were added in the same manner used in FP experiments. Fluorometer setup had donor excitation wavelength at 532 nm, donor emission wavelength at 567 nm, and an acceptor emission wavelength at 668 nm, and 6 nm slit widths. FRET efficiency (E) was calculated by the donor (D) and acceptor (A) emission fluorescence:

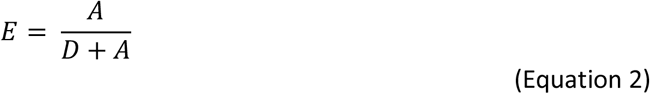

For the BiP_SBD_ FRET assay, a previously described BiP double mutant G518C and Y636C was simultaneously labeled with donor and acceptor fluorophores, AlexaFluor 555 C_2_ maleimide and AlexaFluor 647 C_2_ maleimide, respectively^21^. In the BiP_SBD_ FRET assay, the value of the FRET efficiency is therefore limited to a value of 0.5 because at most only 50% of the BiP molecules can be labeled with one donor and one acceptor fluorophore. BiP was diluted to 50 nM into buffer containing 25 mM MES pH 6.0, 50/150/300/450 mM KCl, 1 mM MgCl_2_, 1 mM ATP, 0.5 mg/mL BSA, and 1 mM DTT. Experiments with ADP contained 1 mM ADP. If indicated, hexokinase-treated ADP was incubated as above experiments. Experiments with 5 % ATP had 1 mM ADP and 0.05 mM ATP and were completed at 50 mM KCl. Fluorometer setup had a donor excitation wavelength at 532 nm, donor emission wavelength at 567 nm, and acceptor emission wavelength at 668 nm, 4 nm slit widths, and 0.5 second integration time. *K*_*D*_ values were calculated using a single-site binding equation,

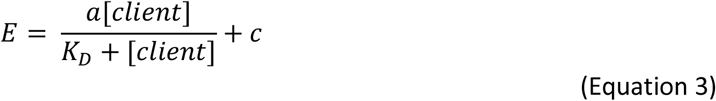

where *a* is the FRET efficiency amplitude and *c* is the FRET efficiency value in the absence of client. *K*_*D*_ values < 0.2 μM were determined via

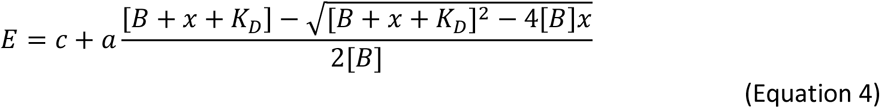

where *x* is the concentration of client E-peptide, *K*_*D*_ is the dissociation constant between BiP and client, and *B* is the concentration of BiP_SBD_ FRET-labeled protein used in experiments.

### Light scattering

ProIGF2 and E-peptide were assayed for light scattering with an absorbance at 320 nm at 25°C. Background was subtracted at 700 nm. 2 μM of each protein was prepared in buffer containing 25 mM MES pH 6.0, 1 mM MgCl_2_, 1 mM ATP, 0.3 mM TCEP. Immediately before assay, proIGF2 was diluted out of urea into buffer containing 50 mM MES pH 6 and 2 mM TCEP.

### Dynamic light scattering

DLS data was obtained using a DLS/SLS-5022F from ALV (ALV-Laser Vertriebsgesellschaft m.b.H.) coupled with a 22mW HeNe Laser from JDS Uniphase Corporation. E-peptide and proIGF2 were diluted into 50 mM MES pH 6.0, 50 mM KCl, 2 mM TCEP, 1 mM MgCl_2_, and 1 mM ATP. 10 rounds of 20 seconds were used for data collection at 25°C. Protein sample was monitored at 90° by laser light scattering at 630 nm. Size-distribution analysis from an intensity correlation function was used to attain R_H_^38^.

### ATPase assay

ATPase activity was measured by depletion of NADH via an enzyme-linked assay with pyruvate kinase and lactose dehydrogenase. 2 μM BiP was assayed in 25 mM MES pH 6.0, 50 mM KCl, 1 mM MgCl_2_, 1 mM ATP, 1 mM DTT, 0.5 mM NADH, 0.5 mM PEP, 0.1 μM pyruvate kinase, 0.1 μM lactose dehydrogenase, and 2.5 μM client protein at 37°C. NADH depletion was monitored at an absorbance of 340 nm. ATPase rates reported are an average of 3 measurements, and the error is the SEM.

## Acknowledgements

We thank Linda Hendershot, Daniel Oprian, and Tijana Ivanovic for providing helpful feedback on the results. Research for this project was supported by NIH R01 GM115356.

## Supplemental Figures & Tables

**Supplemental Figure 1.**
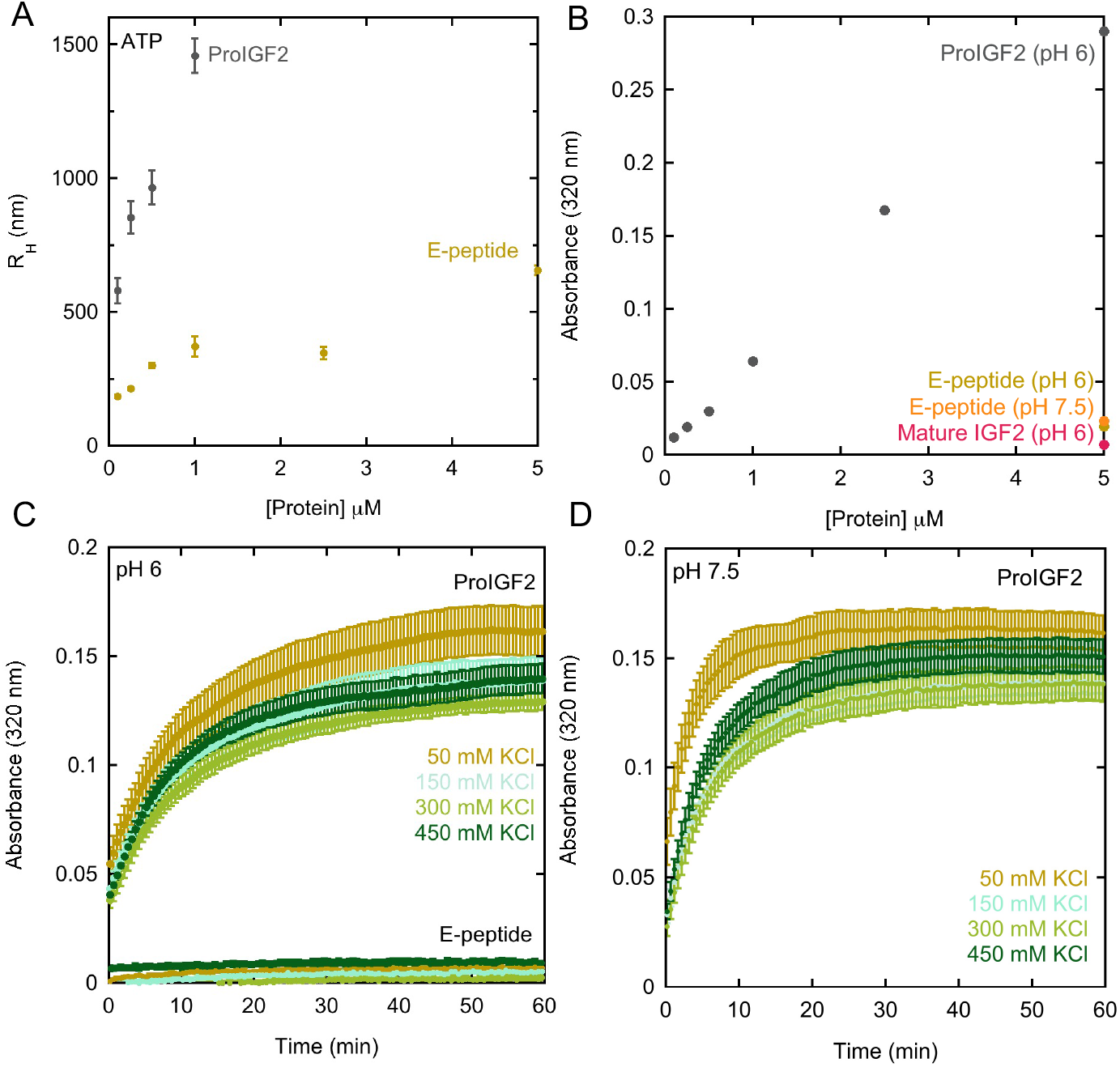
ProIGF2, E-peptide, and mature IGF2 light-scattering (LS) data. **(A)** Average hydrodynamic radius (R_H_) of proIGF2 and E-peptide oligomers as measured by DLS. **(B)** Concentration-dependent LS data with proIGF2 shown with 5 μM E-peptide and mature IGF2. **(C)** LS data with 2 μM proIGF2 and E-peptide at pH 6 and increasing salt concentrations. Rates of proIGF2 LS at pH 6 with increasing KCl are 0.072 ± 0.001, 0.076 ± 0.001, 0.080 ± 0.003, and 0.091 ± 0.001 min^-1^. **(D)** LS data with proIGF2 at pH 7.5 and increasing salt concentrations. Rates of proIGF2 LS at pH 7.5 with increasing KCl are 0.23 ± 0.02, 0.12 ± 0.001, 0.12 ± 0.004, and 0.14 ± 0.004 min^-1^. Absorbance data was collected at 320 nm with a background subtraction of 700 nm. Error bars indicate SEM for 3 replicates.

**Supplemental Figure 2.**
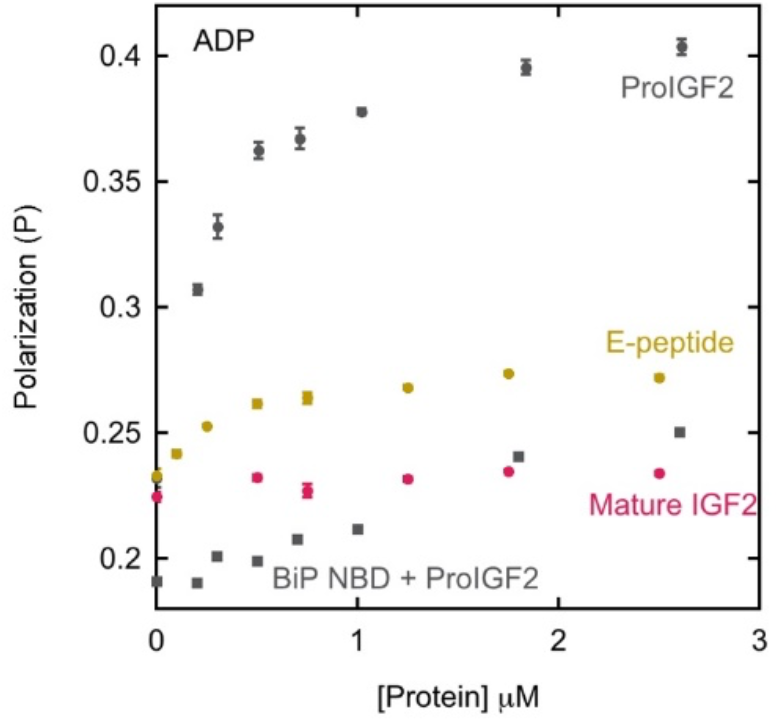
FP data with FITC-BiP and proIGF2, E-peptide and mature IGF2 (circles) and FITC-BiP NBD binding proIGF2 (squares) in ADP conditions.

**Supplemental Figure 3.**
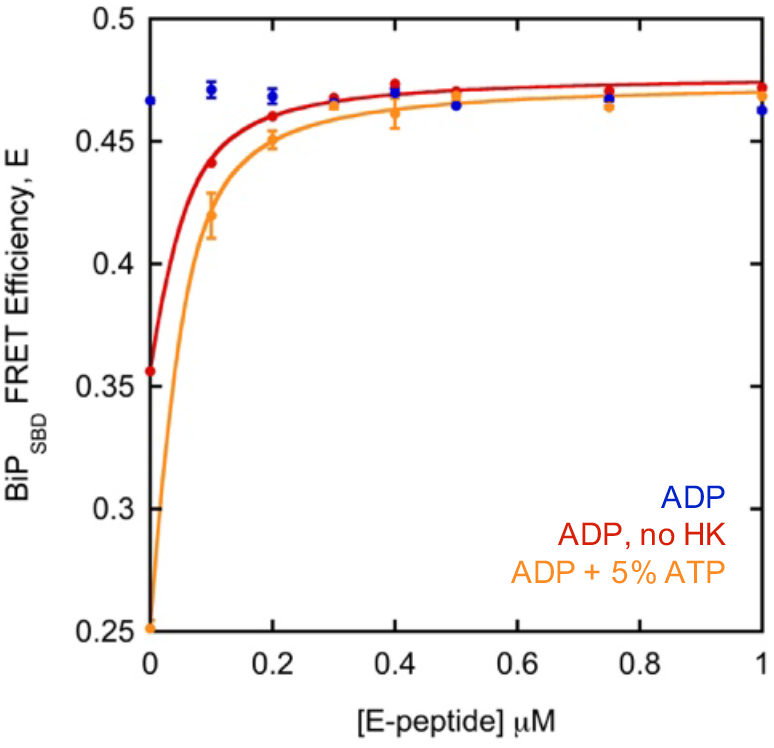
BiP_SBD_ FRET efficiency in the presence of E-peptide with HK-treated ADP (blue), ADP without an HK treatment (red, K_D_ = 0.015 ± 0.005 μM) and ADP + 5% ATP (yellow, K_D_ = 0.019 ± 0.004 μM). Solid line is fit to equation 4. Error bars indicate SEM for at least 3 replicates.

**Supplemental Figure 4.**
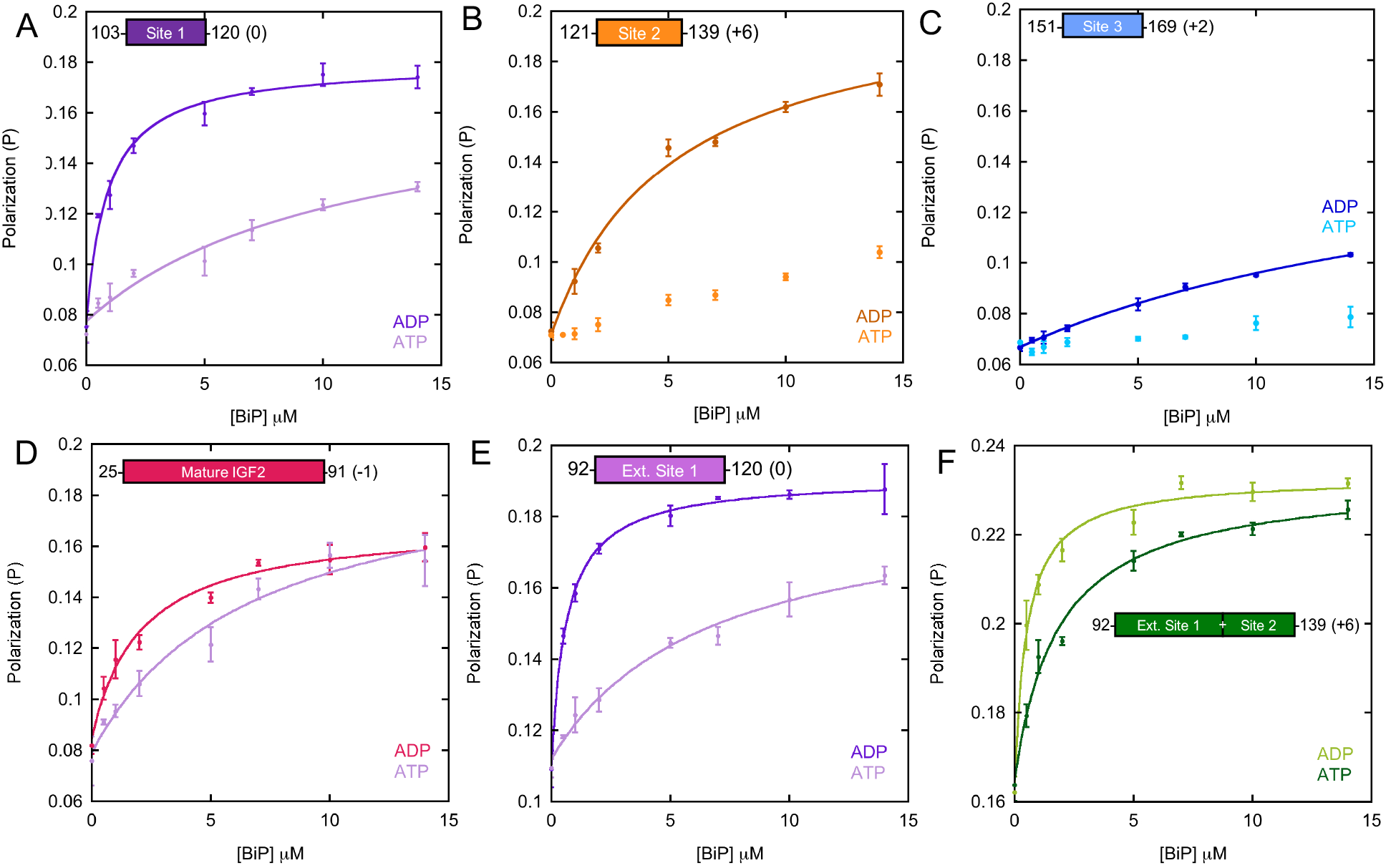
FP BiP binding assay with FITC-labeled E-peptide fragments. Solid lines indicate fit to equation 1, in cases where K_D_ values can be determined. **(A)** BiP affinity for site 1 is: 0.95 ± 0.17 μM and 13 ± 2 μM, under ADP and ATP conditions respectively. K_D_ under ATP conditions is determined using same maximum amplitude as ADP condition. **(B)** BiP affinity for site 2 is 5.3 ± 0.6 μM under ADP conditions. **(C)** BiP affinity for site 3 is 15 ± 1 μM under ADP conditions. **(D)** FP binding assay with FITC-labeled mature IGF2^1 Cys^ and BiP in ADP and ATP states. Fit values as calculated from Equation 1 in the presence of ADP and ATP are 2.7 ± 1.0 and 9.7 ± 4.5 μM, respectively. **(E)** BiP affinity for extended site 1 (residues 92-120) is: 0.67 ± 0.17 μM and 8.0 ± 3.8 μM, under ADP and ATP conditions respectively. **(F)** BiP affinity for extended site 1&2 (residues 92-139) is: 0.52 ± 0.13 μM and 1.9 ± 0.6 μM, under ADP and ATP conditions respectively. All error bars indicate SEM for three replicates.

**Supplemental Figure 5.**
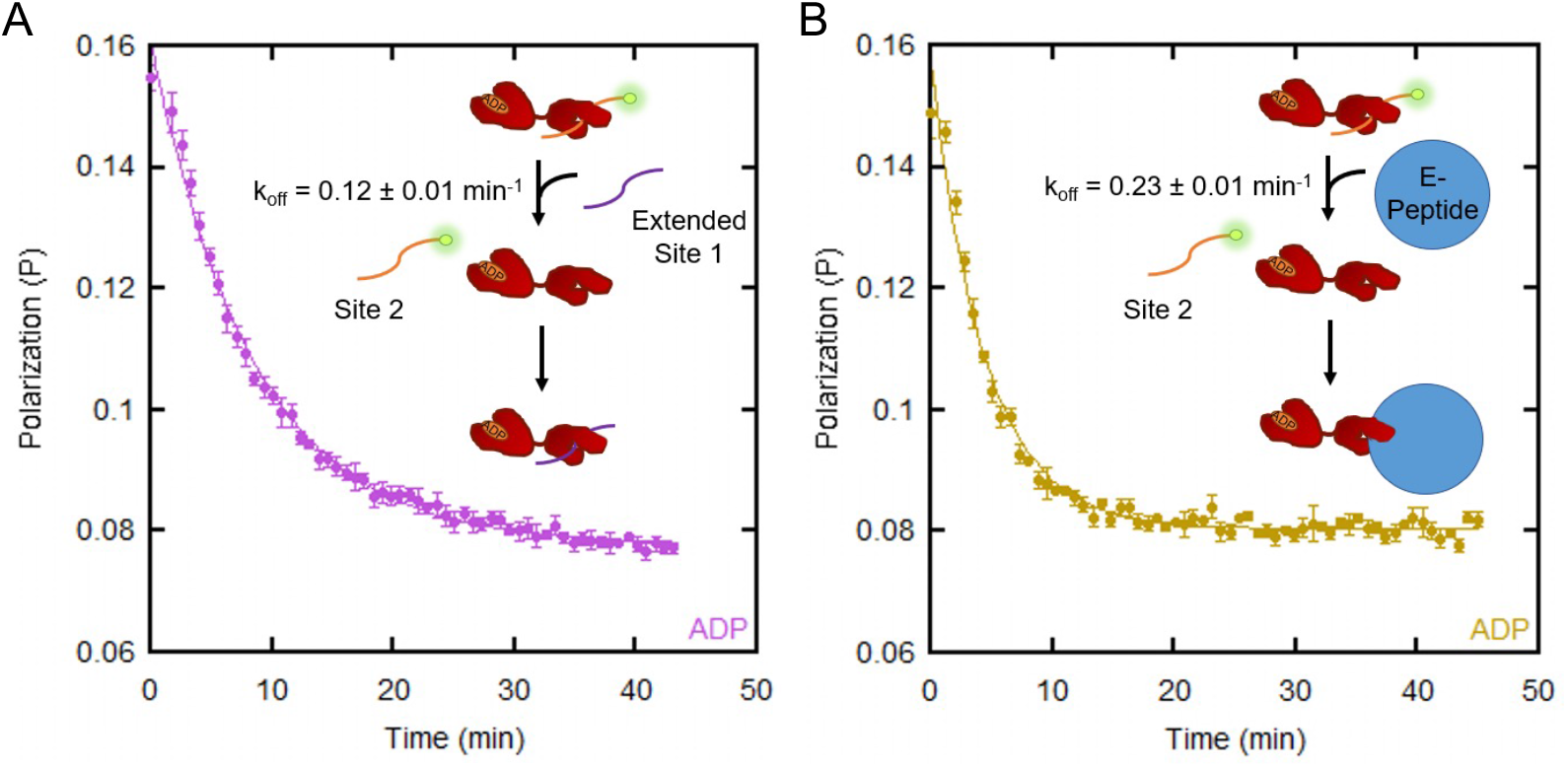
Binding competition experiments. 50 nM FITC-labeled site 2 was prebound to 5.3μM BiP under ADP conditions. **(A)** Competition with 10μM of extended site 1. **(B)** Competition with 5.3μM of oligomerized E-peptide. Solid lines are a fit to an exponential decay. Error bars are the SEM for three replicates.

**Supplemental Figure 6.**
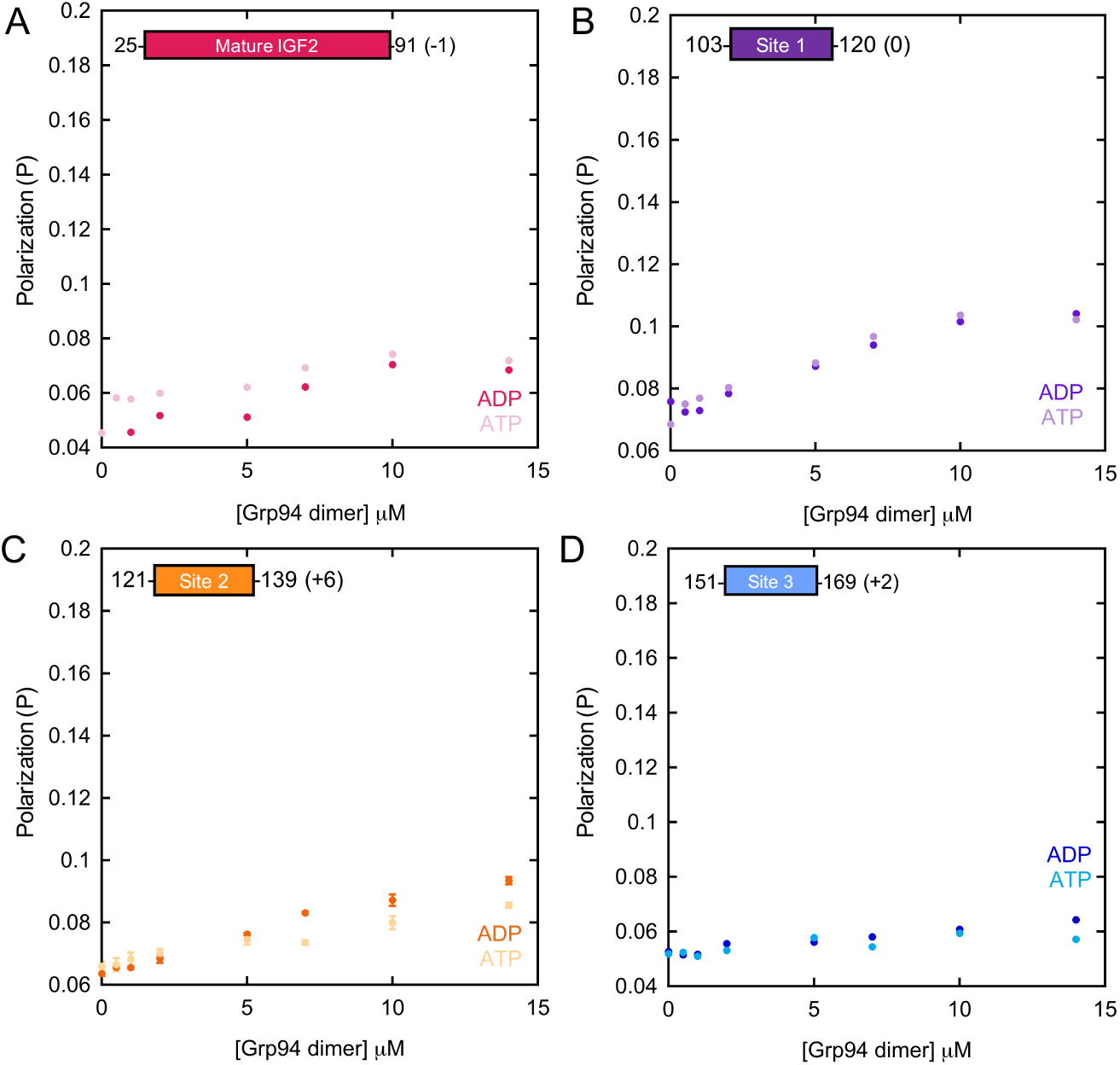
FP assay with FITC-labeled BiP binding-site peptides and mature IGF2 (1 cysteine mutant) in the presence of Grp94 and ADP or ATP. Error bars indicate SEM for 3 replicates, where present. Y-axis is identical to Supplemental Figure 4 with BiP for comparison.

**Supplemental Figure 7.**
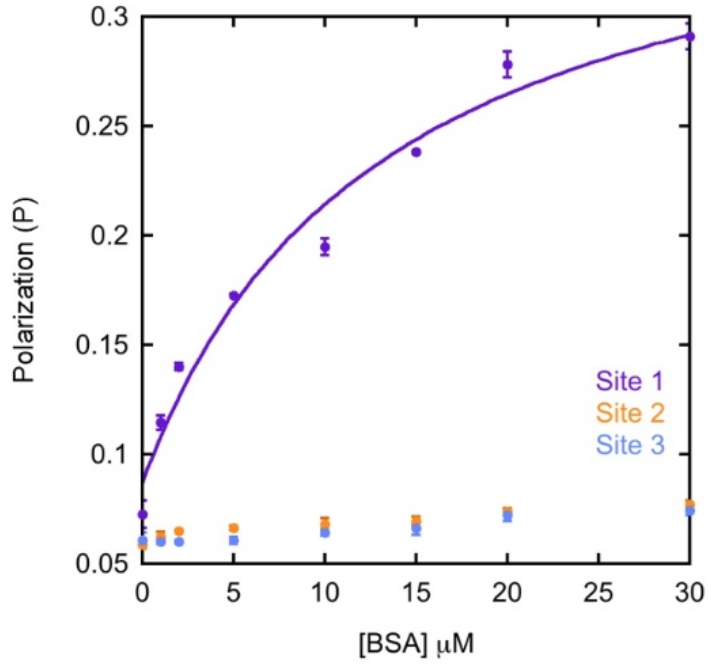
FP assay with FITC-labeled site 1, 2, or 3 binding BSA. Solid line indicates fit to equation 1 (K_D_: 13 ± 2 μM). Error bars indicate SEM of 3 trials.

**Supplemental Figure 8.**
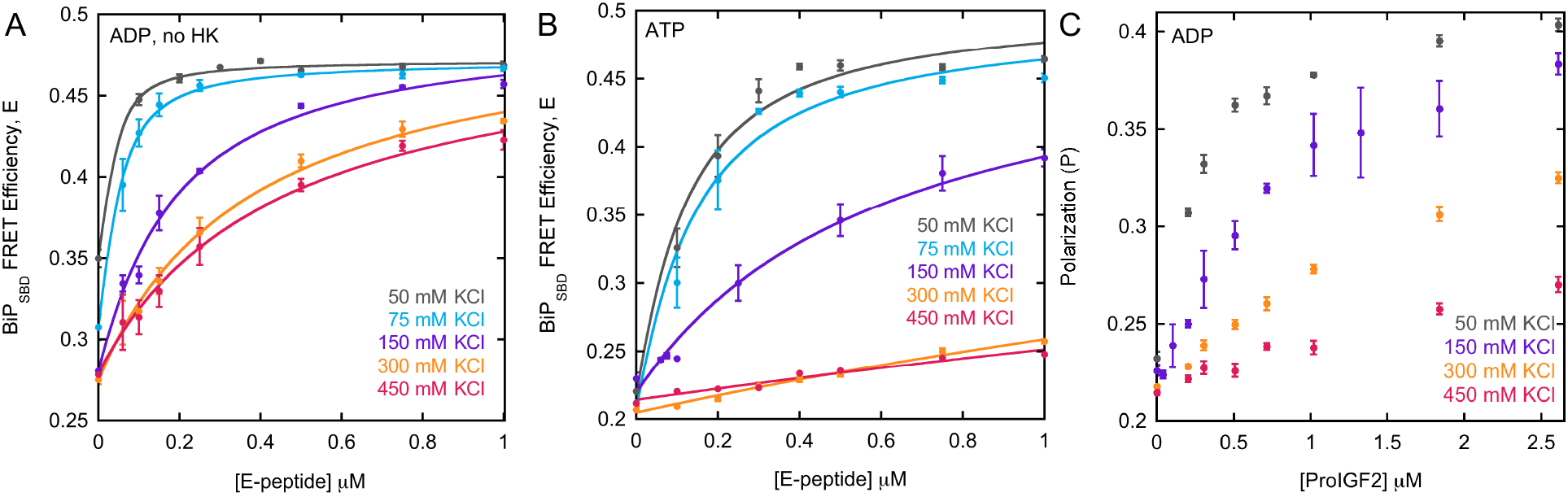
BiP binding E-peptide and proIGF2 at increasing salt concentrations. **(A)** Salt-dependence of BiP_SBD_ FRET assay with E-peptide oligomers and non-HK treated ADP. Fit values of binding affinity are shown in Figure 4A. Solid lines for 50, 75, and 150 mM KCl are a fit to equation 4, and lines for 300 and 450 mM KCl are a fit to equation 3. **(B)** Salt-dependence of the BiP_SBD_ FRET assay with E-peptide oligomers under ATP conditions. Fit values of binding affinity are shown in Figure 4A. Solid lines for 50 and 75 mM KCl are a fit with equation 4, and 150, 300, and 450 mM KCl data are fit with equation 3 and a maximum FRET efficiency set to 0.5. **(C)** BiP binding proIGF2 oligomers as a function of increasing KCl concentration using FP assay with FITC-labeled BiP, in the presence of ADP. Error bars indicate SEM for at least 3 replicates.

**Supplemental Figure 9.**
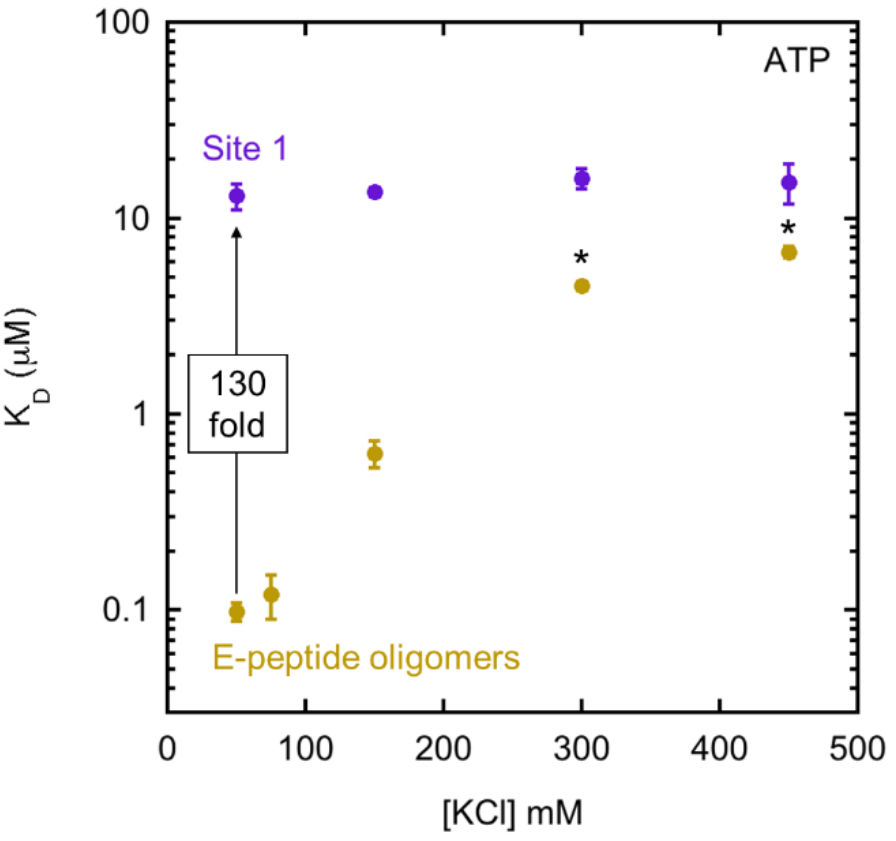
Influence of salt on BiP’s affinities for E-peptide oligomers and site 1 under ATP conditions. K_D_ data from E-peptide 103-120 is from FP data in Supplemental Figure 4A. K_D_ data from E-peptide from Supplemental Figure 8B. Error bars are the SEM for at least three replicates. Asterisks indicate lower confidence of fitting, as described in Supplemental Figure 8B.

**Supplemental Table 1.** Compilation of Hsp70 family dissociation constants towards client proteins. *Oligomerization state for MAPT 3R and 4R not stated. ^#^ Clathrin sequence from *B. taurus* and net charge calculated with one heavy chain and one light chain A. Net charge calculated with one heavy chain and one light chain B is −58. ^@^ Peptide from *C. livia* sequence. ^ indicates synthetic client sequences. All other client sequences are from *H. sapiens*. Peptide clients are assumed to be monomeric.

| Client | Hsp70 | K<sub>d</sub> (μM) | Monomer? | Net charge | Nucleotide | Ref |
| --- | --- | --- | --- | --- | --- | --- |
| C<sub>H</sub>1 domain | BiP | 4.2±0.4 | Yes | +1 | ADP | <sup>31</sup> |
| HTFPAVL peptide | BiP | 11.6±0.6 | Yes | 0 | ADP | <sup>21</sup> |
| MAPT MBD<br>Fibril | Hsp70 | 0.02±0.01 | No | +10 | none | <sup>7</sup> |
| MAPT MBD 10mer+ | Hsp70 | 0.17±0.04 | No | +10 | none | <sup>7</sup> |
| MAPT MBD 6-10mer | Hsp70 | 0.34±0.09 | No | +10 | none | <sup>7</sup> |
| MAPT MBD 3-5mer | Hsp70 | 0.97±0.19 | No | +10 | none | <sup>7</sup> |
| MAPT MBD 1-2mer | Hsp70 | 5.6±1.4 | Yes | +10 | none | <sup>7</sup> |
| MAPT 3R* | Hsc70 | 0.31±0.05 | * | +10 | none | <sup>36</sup> |
| MAPT 4R* | Hsc70 | 0.16±0.04 | * | +13 | none | <sup>36</sup> |
| α-synuclein fibril | Hsp70 | 5.8±0.4 | No | -9 | ATP | <sup>8</sup> |
| α-synuclein monomer | Hsp70 | ~10 | Yes | -9 | ATP | <sup>8</sup> |
| NR peptide^ | BiP | 0.95 | Yes | +1 | none | <sup>39</sup> |
| leukocyte antigen B*2702-derived peptide Bw4 | Hsp70 | 1.8 | Yes | +2 | ATP | <sup>40</sup> |
| NLLRLTGW^ (Javelin 1) | Hsp70 | 0.9 | Yes | +1 | ADP | <sup>34</sup> |
| Faf1 peptide^ (FYQLALT) | Hsc70 | 4.3±0.9 | Yes | 0 | ADP | <sup>25</sup> |
| Faf1 peptide^ (FYQLALT) | Hsc70 | 37-51 | Yes | 0 | ATP | <sup>25</sup> |
| Clathrin<sup>#</sup> | Hsp70 | 3 | No | -64 | 90% ADP,<br>10% ATP | <sup>26</sup> |
| Clathrin<sup>#</sup> | Hsp70 | 12 | No | -64 | ATP | <sup>26</sup> |
| Cytochrome c peptide<sup>@</sup><br>(IFAGIKKKAERADLIAYLKQAT<br>AK) | Hsp70 | 7 | Yes | +4 | 90% ADP,<br>10% ATP | <sup>26</sup> |
| Cytochrome c peptide<sup>@</sup> | Hsp70 | 300 | Yes | +4 | ATP | <sup>26</sup> |

**Supplemental Table 2.** Compilation of representative DnaK dissociation constants towards client proteins. *CALLQSRLLLSAPRRAAATARA, derivative of chicken mitochondrial aspartate aminotransferase signal sequence.

| Client | K<sub>d</sub> (μM) | Monomer? | Net charge | Nucleotide | Ref |
| --- | --- | --- | --- | --- | --- |
| Peptide pp* | 0.06 | Yes | +4 | ADP | <sup>41</sup> |
| Peptide pp* | 2.2 | Yes | +4 | ATP | <sup>41</sup> |
| human telomere repeat binding factor 1 (hTRF1) 377-430 | 1.4±0.2 | Yes | +10 | ADP | <sup>42</sup> |
| human telomere repeat binding factor 1 (hTRF1) 377-430 | 18±3 | Yes | +10 | ATP | <sup>42</sup> |
| Islet amyloid polypeptide (*H. sapiens*) | ~pM | No | +2 | apo | <sup>35</sup> |
| σ<sup>32</sup> peptide (Q132-Q144) (*E. coli*) | 0.078 | Yes | +5 | apo | <sup>43</sup> |
| σ<sup>32</sup> (*E. coli*) | 5 | Yes | -6 | apo | <sup>44</sup> |

